# GLASS: assisted and standardized assessment of gene variations from Sanger sequence trace data

**DOI:** 10.1101/088401

**Authors:** Karol Pal, Vojtech Bystry, Tomas Reigl, Martin Demko, Adam Krejci, Tasoula Touloumenidou, Evangelia Stalika, Boris Tichy, Paolo Ghia, Kostas Stamatopoulos, Sarka Pospisilova, Jitka Malcikova, Nikos Darzentas, European Research Initiative on CLL (ERIC) – TP53 Network

## Abstract

**Motivation:** Sanger sequencing remains the reference method for sequence variant detection, especially in a clinical setting. However, chromatogram interpretation often requires manual inspection and in some cases considerable expertise. Additionally, variant reporting and nomenclature is typically left to the user, which can lead to inconsistencies.

**Results:** We introduce GLASS, a tool built to assist with the assessment of gene variations in Sanger sequencing data. Critically, it provides a standardized variant output as recommended by the Human Genome Variation Society.

**Availability:** The program is freely available online at <u>http://bat.infspire.org/genomepd/glass/</u>.

**Supplementary information:** Supplementary data are available at Bioinformatics online.

## 1 Introduction

Despite great advances in next-generation sequencing, Sanger sequencing still represents the ‘golden standard’ for variant detection and validation, in both research and clinical applications. Moreover, for the majority of diagnostic laboratories, it remains the most applicable approach considering cost and number of samples analyzed.

The analysis of Sanger sequence trace data is also still a laborious task and, most importantly, it can be compromised by a lack of expertise and experience. Several software tools exist to ease this analysis, and these can be helpful but also detrimental. Naïve trace viewers for manual inspection can be cumbersome and ultimately error-prone, while commercial solutions can be prohibitively expensive and their advanced features may prove to be too complex and even discouraging. More often than not, different users use different solutions, resulting in further complications with regards to the consistency of the reported results. Such conclusions were confirmed within the TP53 Network of the European Research Initiative on CLL / ERIC (ericll.org/pages/networks/tp53network) through a survey of the participants of the inaugural Workshop on TP53 mutation analysis (ericll.org/pages/meeting/1st-eric-workshop-on-tp53-analysis-in-chroniclymphocytic-leukemia).

In this context, we developed and herein describe GLASS, which provides an intuitive way for both novice and experienced practitioners to discover and assess gene variations from Sanger sequence trace data. Importantly, the application reports sequence variants in a standardized way as recommended by the Human Genome Variation Society (HGVS, varnomen.hgvs.org), so that users can be valid and consistent in their clinical and research activities and collaborations.

## 2 Methods

### Implementation

GLASS is written in R and uses Shiny (Chang *et al.*, 2016, R package) for client-server interactions and interface rendering. A custom D3-based (Bostock *et al.*, 2011) JavaScript library was developed for rendering chromatograms. GLASS is showcased in Figure 1 and its basic components described below.

**Figure 1.**
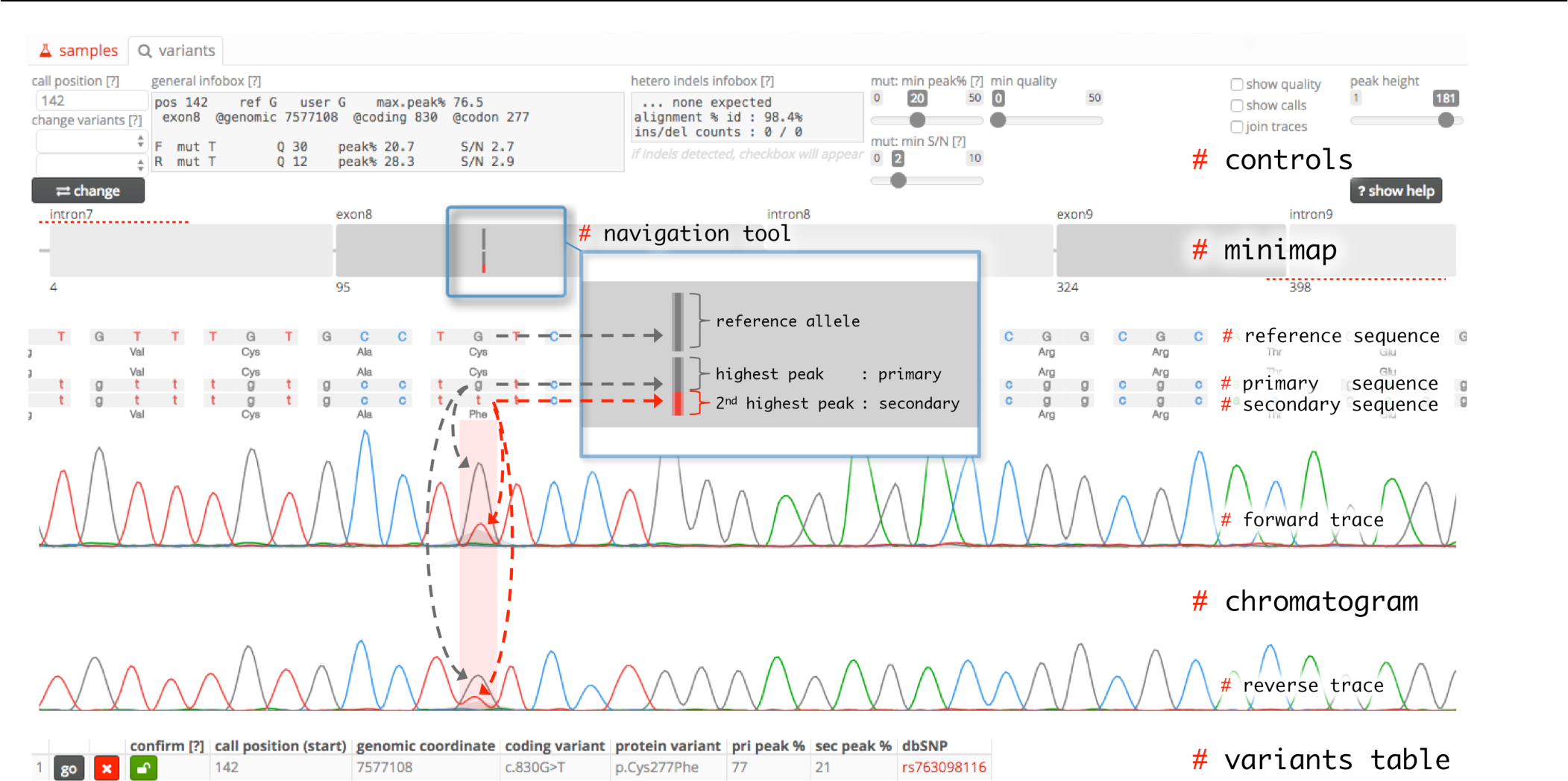
The ‘variants’ panel. Actual screenshot of the ‘variants’ panel, with annotations (#-tagged labels, dashed arrows, zoomed-in inset) manually added for presentation. The ‘minimap’ shows the alignment to the reference gene (in this case a *TP53* gene region spanning introns 7 to 9) and the detected variant. The ‘navigation tool’ (blue rectangle, also as zoomed-in inset) selects the region to display in the ‘chromatogram’. In the ‘chromatogram’ the detected variant is highlighted with a pink vertical bar, and the exportable information is printed in the ‘variants table’. Grey/red dashed arrows added to show the link between the primary/secondary peaks, the primary/secondary sequence, and the minimap variant representation.

### Sequence references

GLASS only uses its own curated reference sequences for standardization and consistency. Users can request the addition of new ones, which requires expert validation. GLASS can also be used without references, acting like a trace viewer.

### Sample file handling

Files are handled in the ‘samples’ panel (supplementary figure 1). GLASS can automatically detect the reference gene and the orientation of the sequence, however users can override these.

### Controls

The ‘variants’ panel features controls for changing base calls and setting parameters, e.g. the sensitivity at which somatic mutations will be detected. The ‘general infobox’ shows information of any position selected in the chromatogram. Information on heterozygous insertions/deletions (indels) is displayed in the ‘hetero indels infobox’.

### Minimap

The ‘minimap’ presents an overview of aligned exons and introns of the reference gene, which can also be highlighted in case of splice variants (supplementary figure 3). Candidate sequence variants appear with an intuitive structure and color code representing the reference base and the primary and secondary peaks (Figure 1 inset).

### Chromatogram

The ‘chromatogram’ plots the traces, with peak widths normalized for better forward and reverse strand alignment. Each peak is annotated with the reference sequence, the primary and secondary sequences, cDNA and genomic coordinates, call qualities, and amino acid translations in the case of exons.

### Variant calling

Germline variants appear as a single peak (homozygous) or as two peaks of lower but similar height (heterozygous) and are easily identifiable. For the more challenging somatic variants, GLASS estimates local background noise and calculates the signal to noise ratio for the second highest peak. If this ratio and the secondary peak (second highest peak) are above user-settable thresholds, the variant is reported.

Homozygous indels are recognized as an alignment gap either in the base calls (deletion) or the reference sequence (insertion). Heterozygous and somatic indels manifest as a series of minor (secondary) peaks after the indel site on the forward strand and before the site on the reverse strand. GLASS can automatically detect the emerging secondary sequences, re-align them and identify the candidate indel, in which case it alerts the user to consider the suggested correction of the alignment (supplementary figure 2).

Additionally, GLASS aligns reference introns and exons separately, which allows for correct identification of alternative splicing, e.g. the *TP53* beta-variant (Flaman *et al.*, 1996) (supplementary figure 3) and also aberrant transcripts. This is especially important for the analysis of FASAY (Flaman *et al.*, 1995) data where whole introns and exons may ‘drop in’ or ‘fall out’.

### Variants table

All candidate variants appear in detail in the variants table, with emphasis on proper nomenclature and presentation according to HGVS. Upon inspection, the user can ‘confirm’ the variant, which will then appear in the ‘samples’ panel and become available for export.

## 3 Results

GLASS is continuously validated against a growing set of manually analyzed samples with expertly confirmed mutations. Most samples are taken directly from the official certification activities of ERIC, with others originating from co-authoring laboratories. As showcased in the Supplementary Material, GLASS was able to identify correctly and name properly all the mutations in the TP53 Analysis Certification set. It is important to note that user interaction is necessary in some cases, since GLASS is built to assist and not automate this process.

## 4 Conclusions

GLASS is a bioinformatic implementation of best practices of labs with published know-how in the analysis of *TP53* and other clinically relevant genes. It was specifically developed in the context of and for the educational and certification activities of the ERIC TP53 Network. We hope and believe it will be useful to researchers and clinical practitioners as an intuitive tool for standardized Sanger trace data analysis.

Future plans include the ability to resume and share analyses; the concatenation of multiple files / amplicons from the same patient to cover the complete reference sequence; the ability to load annotations, e.g. known mutations or hotspots, for visualizations and variant calling.

## Acknowledgements

Computational resources in CEITEC MU were provided by MetaCentrum (LM2010005) and CERIT-SC (CERIT Scientific Cloud, Operational Program Research and Development for Innovations, Reg. no. CZ.1.05/3.2.00/08.0144).

## References

Bostock,M. et al. (2011) D3: Data-Driven Documents. IEEE Trans Vis. Comp Graph. Proc InfoVis.

Flaman,J.M. et al. (1995) A simple p53 functional assay for screening cell lines, blood, and tumors. Proc. Natl. Acad. Sci. U. S. A., 92, 3963–3967.

Flaman,J.M. et al. (1996) The human tumour suppressor gene p53 is alternatively spliced in normal cells. Oncogene, 12, 813–818.

